## Supplementary Information for "Inferring neuron-neuron communications from single-cell transcriptomics through NeuronChat"

This file includes Supplementary Figures 1-13.

### Supplementary Figures

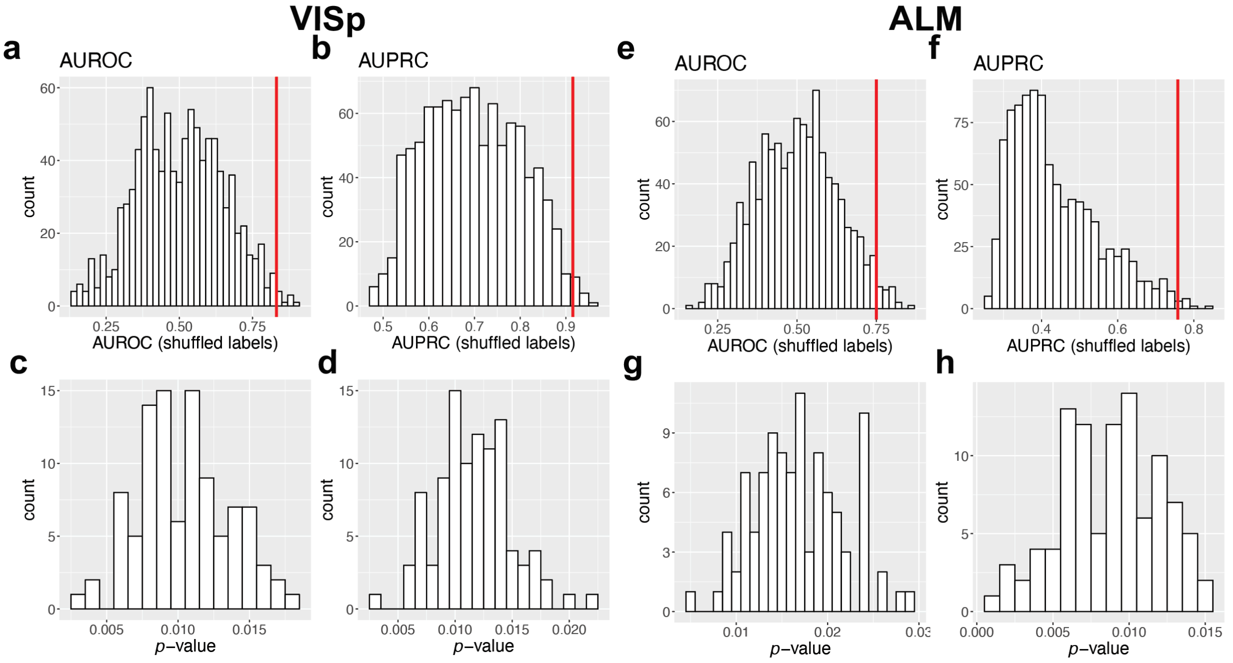


**Supplementary Figure 1. Benchmarking NeuronChat with shuffled ground truth labels.**

(a-b) The distribution of AUROC (a) and AUPRC (b) values for 1,000 times of cell type label shuffling in the VISp projection network. The red line indicates the original AUROC/AUPRC for the VISp projection network without label shuffling.

(c-d) The distribution of *p*-values for AUROC (c) and AUPRC (d). The *p*-value is defined as the proportion of AUROC/AUPRC values that are larger than or equal to the original one (indicated by the red line in a-b), based on 100 independent repeated simulations. Mean±SD for the *p*-values is 0.010±0.0036 for AUROC and 0.012±0.0038 for AUPRC.

(e-h) Repeat analysis for ALM projection network, analogous to (a-d). Mean±SD for the *p*-values is 0.017±0.0048 for AUROC and 0.0087±0.0033 for AUPRC.

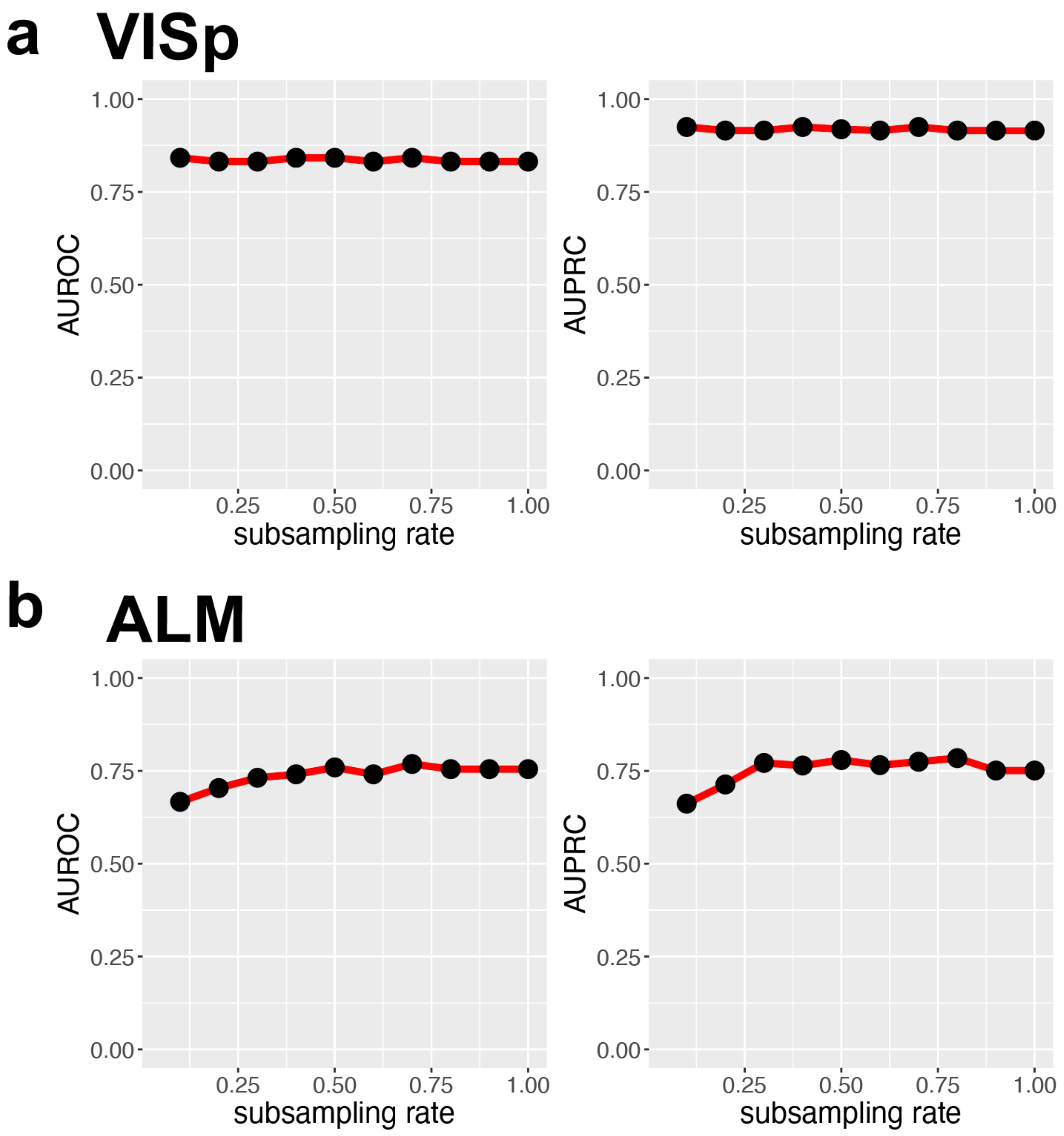


**Supplementary Figure 2. The robustness of NeuronChat to subsampling.**

(a) The AUROC (left panel) and AUPRC (right panel) of the predicted projection network of VISp as a function of subsampling rate.

(b) The AUROC (left panel) and AUPRC (right panel) of the predicted projection network of ALM as a function of subsampling rate.

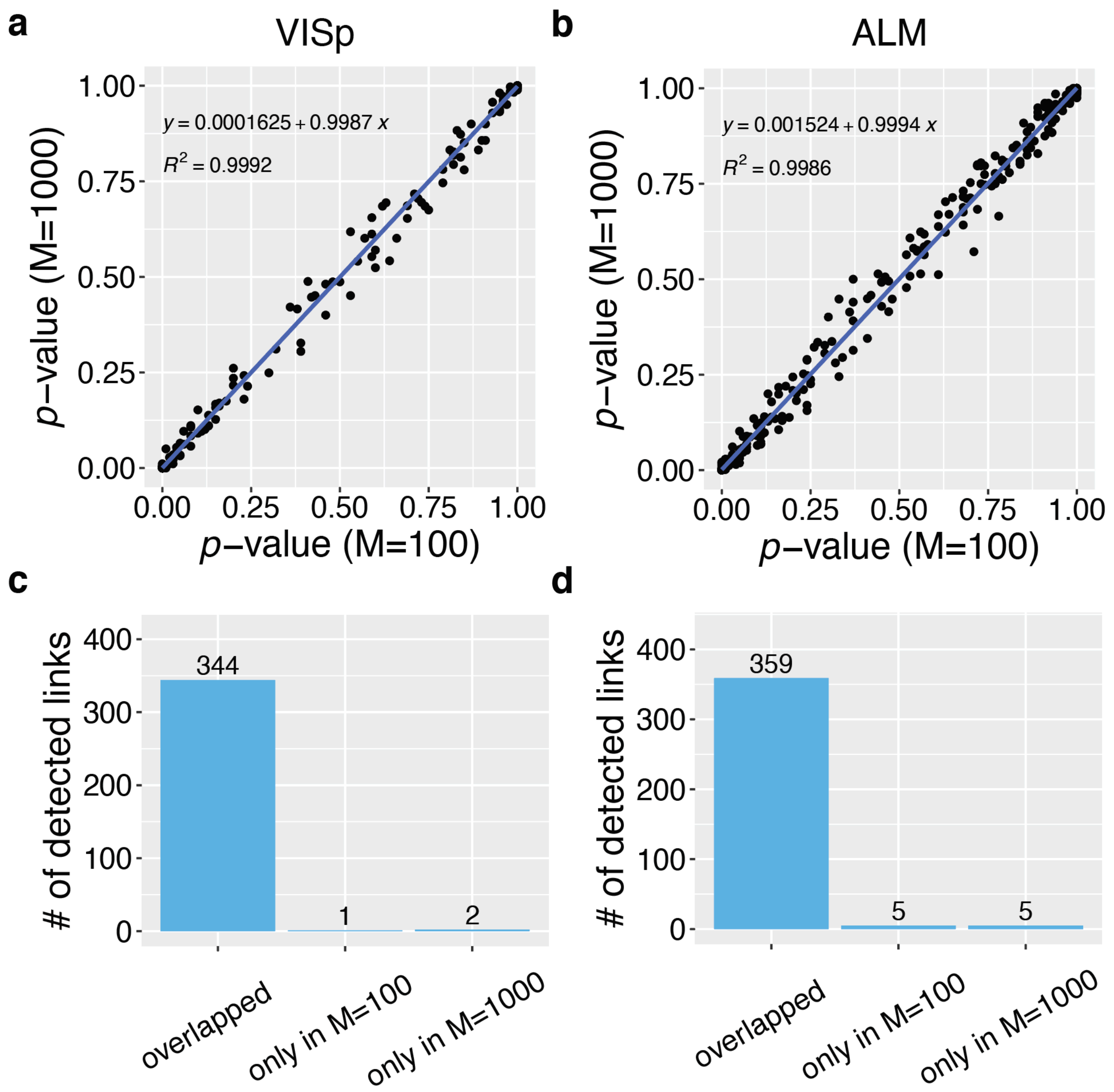


**Supplementary Figure 3. Comparison of *p*-values and number of links calculated by 100 and 1,000 permutations.**

(a-b) The scatter plot of *p*-values calculated by 100 and 1,000 permutations for VISp (a) and ALM (b) projection network. The dots in each plot represents all non-zero cell-cell communication links for all possible interaction pairs. The plot shows the permutation test's original *p*-values that are not adjusted by Benjamini-Hochberg procedure. The regression line, regression equation, and adjusted R-squared are shown in each plot.

(c-d) Comparison of the number of links detected by 100 and 1,000 permutations for VISp (c) and ALM (d) projection network.

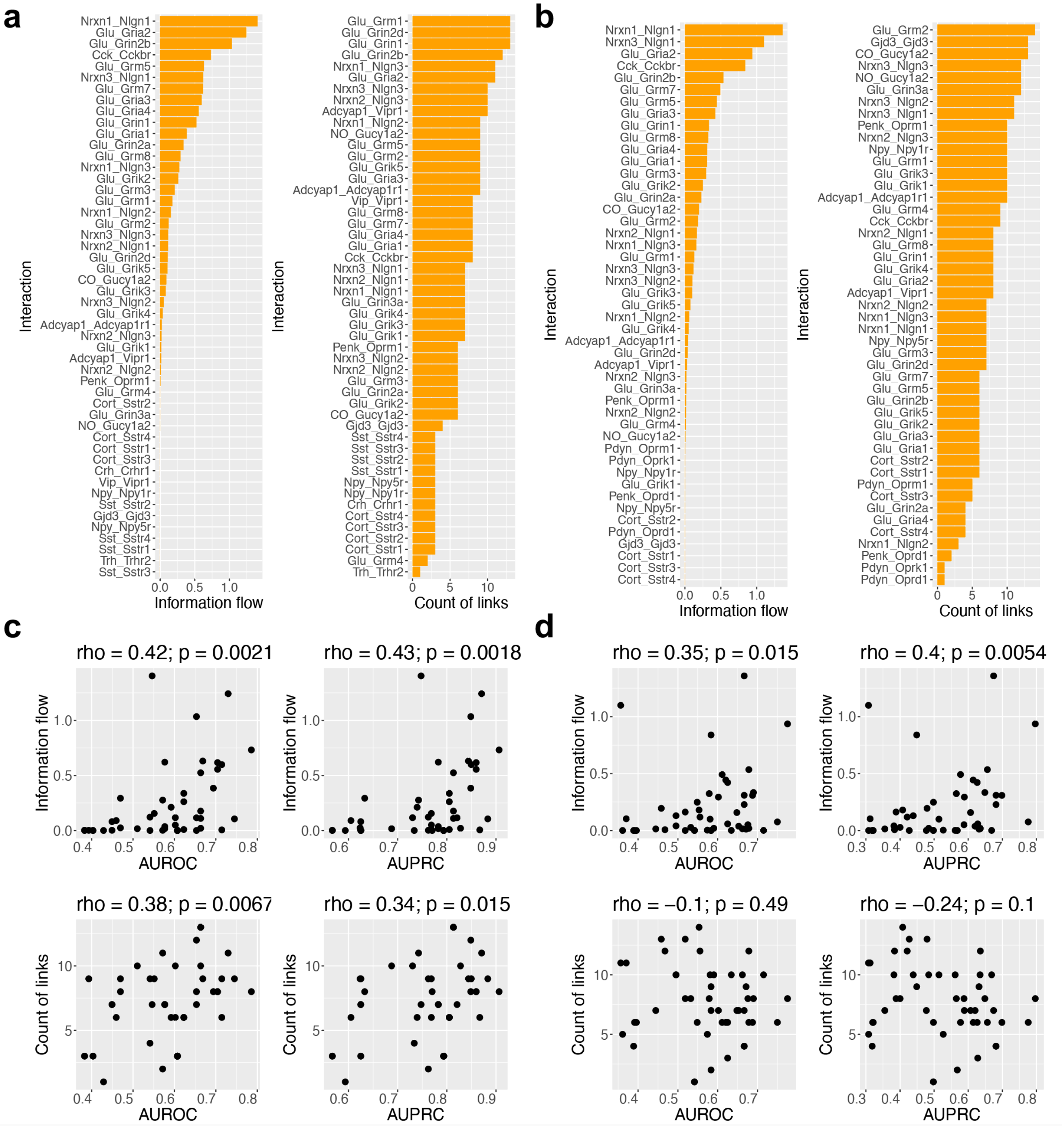


**Supplementary Figure 4. Information flow and count of links for individual interaction pairs and their relationships to the capability of predicting the projection networks of VISp and ALM.**

(a-b) Bar charts illustrating the information flow (left) and the count of links (right) for each interaction pair for VISp (a) and ALM (b). The information flow for a given interaction pair is defined by the sum of communication probability over all significant links among all cell types.

(c-d) Scatter plots of information flow and count of links versus AUROC and AUPRC for VISp (c) and ALM (d). The Spearman's rank correlation coefficient and corresponding *p*-values are indicated on the top of each subplot.

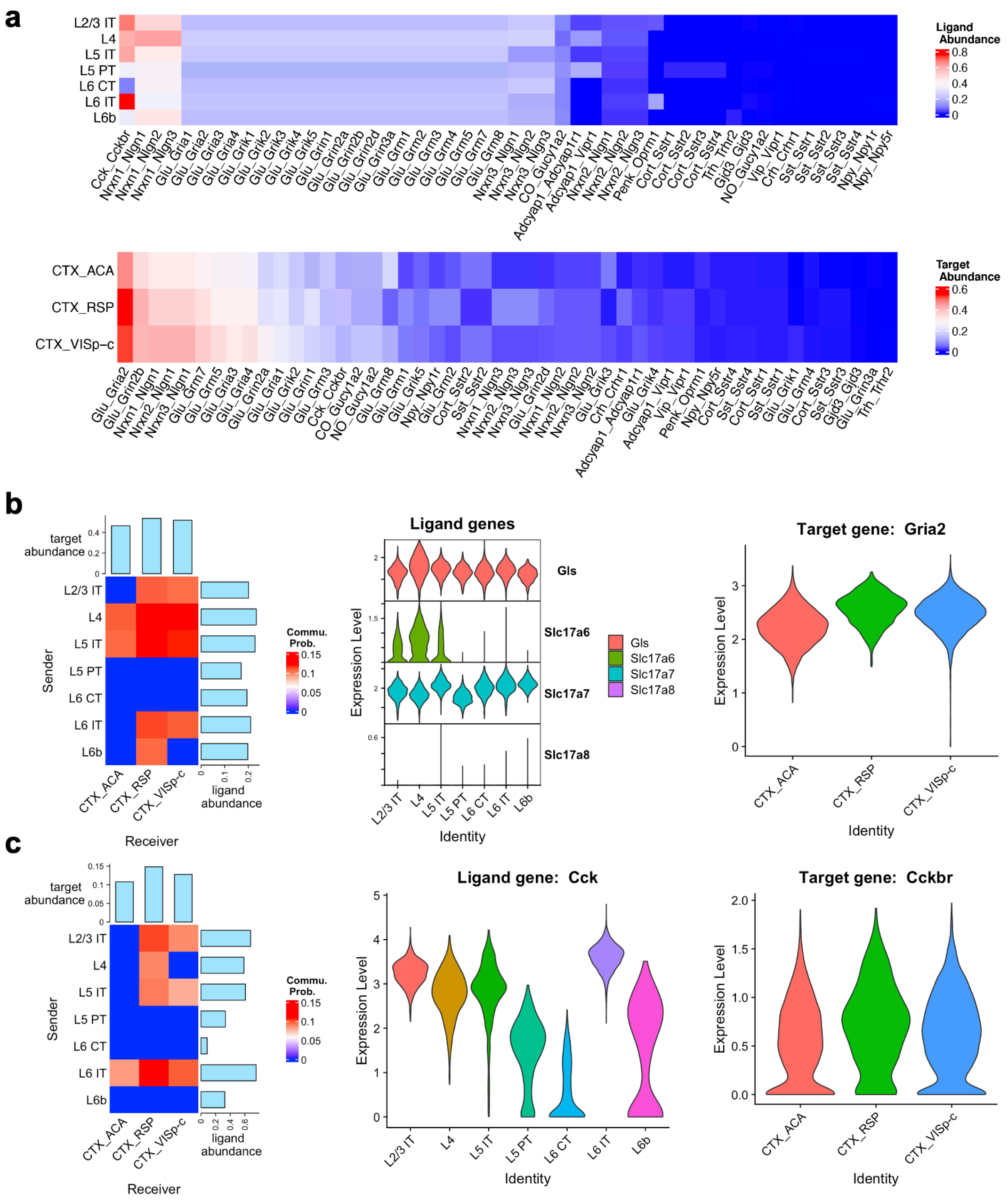

**Supplementary Figure 5. Examples illustrating how variations in the abundance profiles of ligands in sending cell groups and targets in receiving cell groups differentiate the communication strength for VISp projection network.**

(a) Heatmaps of the ligand abundance in the sending cell groups (upper panel) and target abundance in the receiving cell groups (bottom panel) for the interaction pairs with at least one significant link. For each panel, the interaction pairs are sorted by the sum of ligand (or target) abundance over the sending (or receiving) cell groups.

(b-c) The intercellular networks (left panel), the expression of the ligand-related gene(s) (middle panel), and the expression of the target (right panel) for the interaction pairs Glutamate_Gria2 (b) and Cck-Cckbr (c).

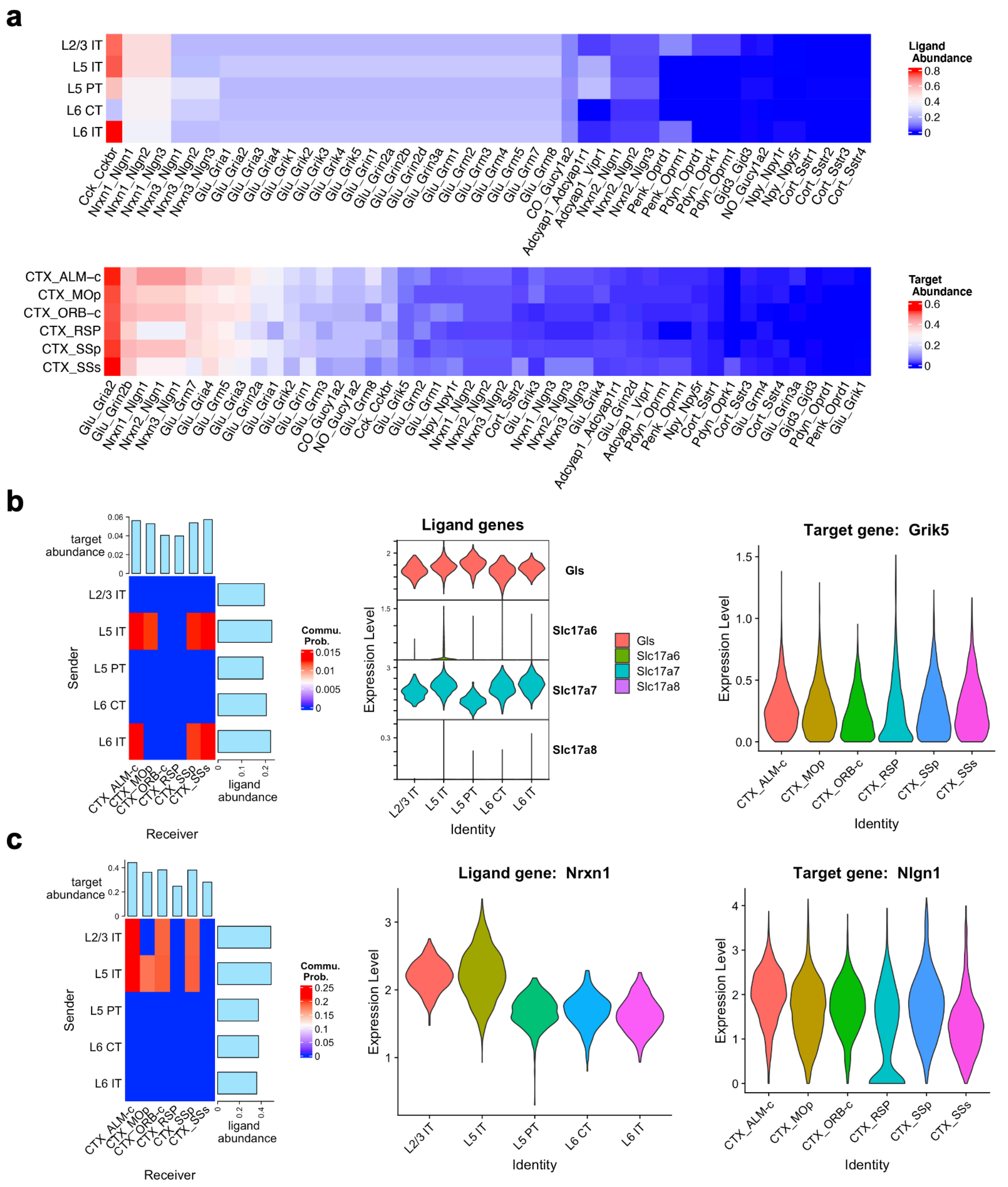


**Supplementary Figure 6. Examples illustrating how variations in the abundance profiles of ligands in sending cell groups and targets in receiving cell groups differentiate the communication strength for ALM projection network.**

(a) Heatmaps of the ligand abundance in the sending cell groups (upper panel) and target abundance in the receiving cell groups (bottom panel) for the interaction pairs with at least one significant link. For each panel, the interaction pairs are sorted by the sum of ligand (or target) abundance over the sending (or receiving) cell groups.

(b-c) The intercellular networks (left panel), the expression of the ligand-related gene(s) (middle panel), and the expression of the target (right panel) for the interaction pairs Glutamate_Grik5 (b) and Nrxn1-Nlgn1 (c).

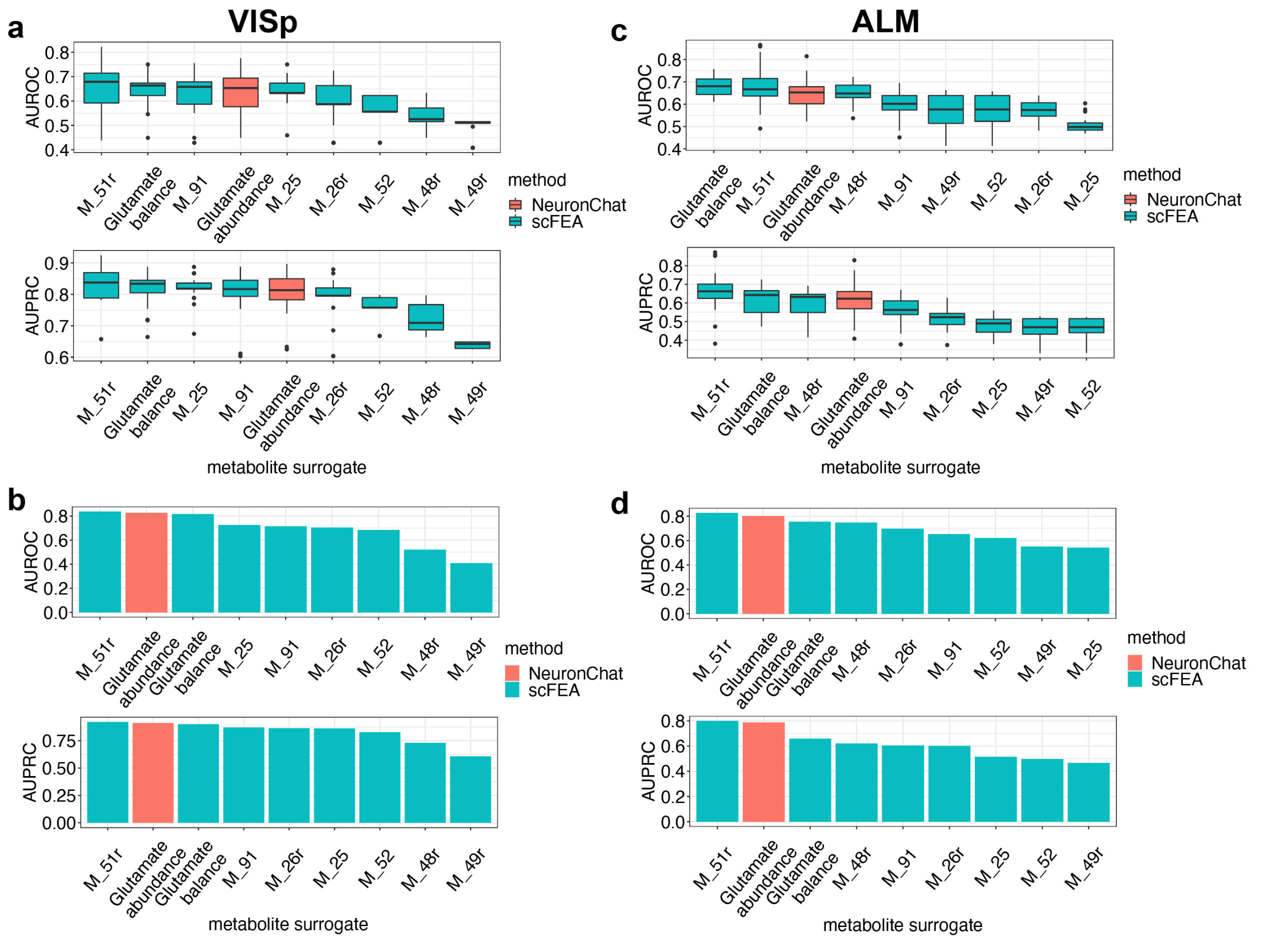


**Supplementary Figure 7. Comparison between NeuronChat's ligand abundance and scFEA-derived metabolite surrogates in identifying glutamate-mediated communication networks.**

(a) Boxplots of AUROC (upper panel) and AUPRC (lower panel) values of 24 glutamate-mediated communication networks for predicting the VISp projection network, for the nine glutamate surrogates. For the metabolic module representing the opposite direction of glutamate accumulation (e.g., M_48), we use the maximum flux value among all cells minus the original flux as the surrogate for glutamate, denoted by the original module name with a suffix "r" (same for b, c, and d). Boxplot elements: center line, median; box limits, upper and lower quartiles; whiskers, 1.5x interquartile range; points, outliers.

(b) Barplots of AUROC (upper panel) and AUPRC (lower panel) values of the aggregated glutamate-mediated communication networks for predicting VISp projection network, for the nine glutamate surrogates.

(c-d) Repeat analysis for ALM projection network, analogous to (a-b).

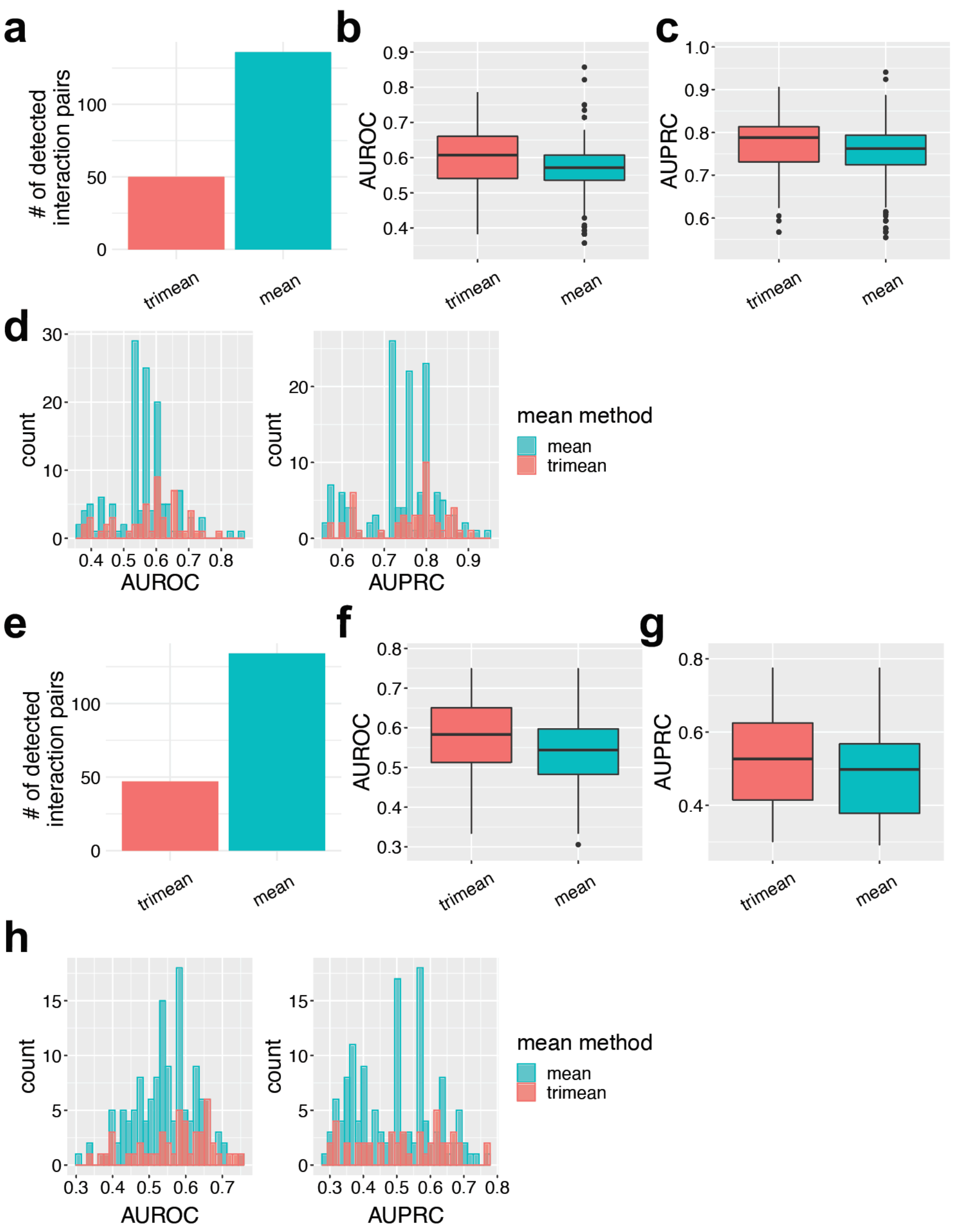


**Supplementary Figure 8. Comparison of Tukey's trimean and arithmetic mean in predicting VISp and ALM projection networks.**

(a) The number of detected interaction pairs for the VISp projection networks for the two mean methods.

(b-c) Boxplots showing the AUROC (b) and AUPRC (c) values of the individual VISp projection networks for the two mean methods. Boxplot elements: center line, median; box limits, upper and lower quartiles; whiskers, 1.5x interquartile range; points, outliers.

(d) Distributions of AUROC (left panel) and AUPRC (right panel) values of the individual VISp projection networks for the two mean methods.

(e-h) Repeat analysis for ALM projection network, analogous to (a-d).

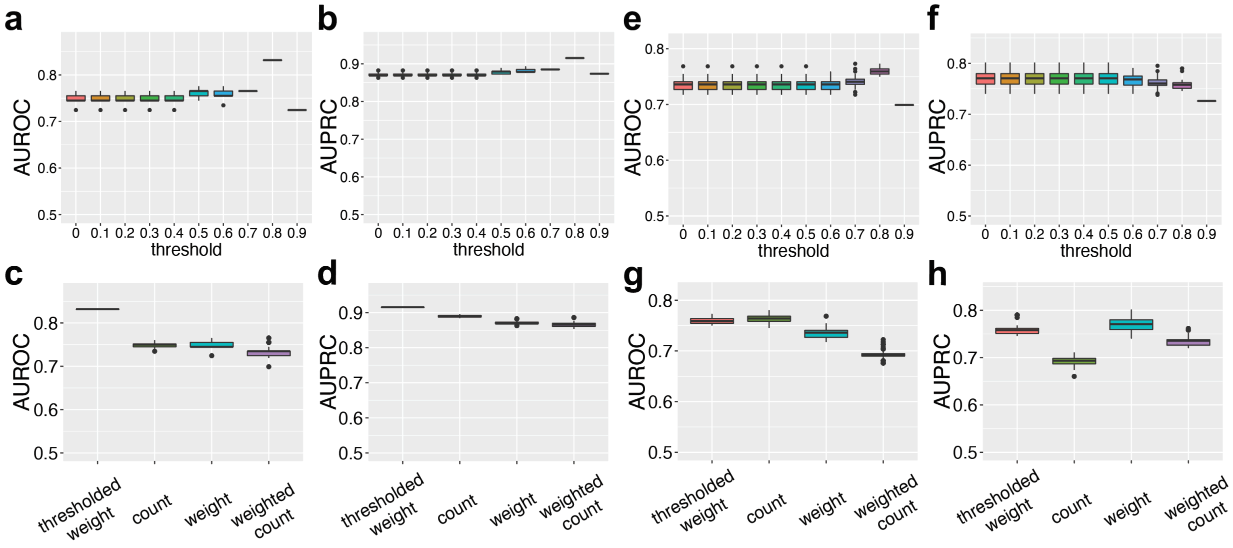


**Supplementary Figure 9. Optimization of the aggregation method.**

(a-b) Boxplots of AUROC (a) and AUPRC (b) values for 100 repeats of the aggregated VISp projection networks at different quantile thresholds of the aggregation method “thresholded weight”. Boxplot elements: center line, median; box limits, upper and lower quartiles; whiskers, 1.5x interquartile range; points, outliers.

(c-d) Boxplots of AUROC (c) and AUPRC (d) values for 100 repeats of the aggregated VISp projection networks for four different aggregation methods. For the aggregation method “thresholded weight”, the thresholding parameter is chosen as 80% quantile of communication strength values for each interaction pair. Boxplot elements: center line, median; box limits, upper and lower quartiles; whiskers, 1.5x interquartile range; points, outliers.

(e-h) Repeat analysis for ALM projection network, analogous to (a-d).

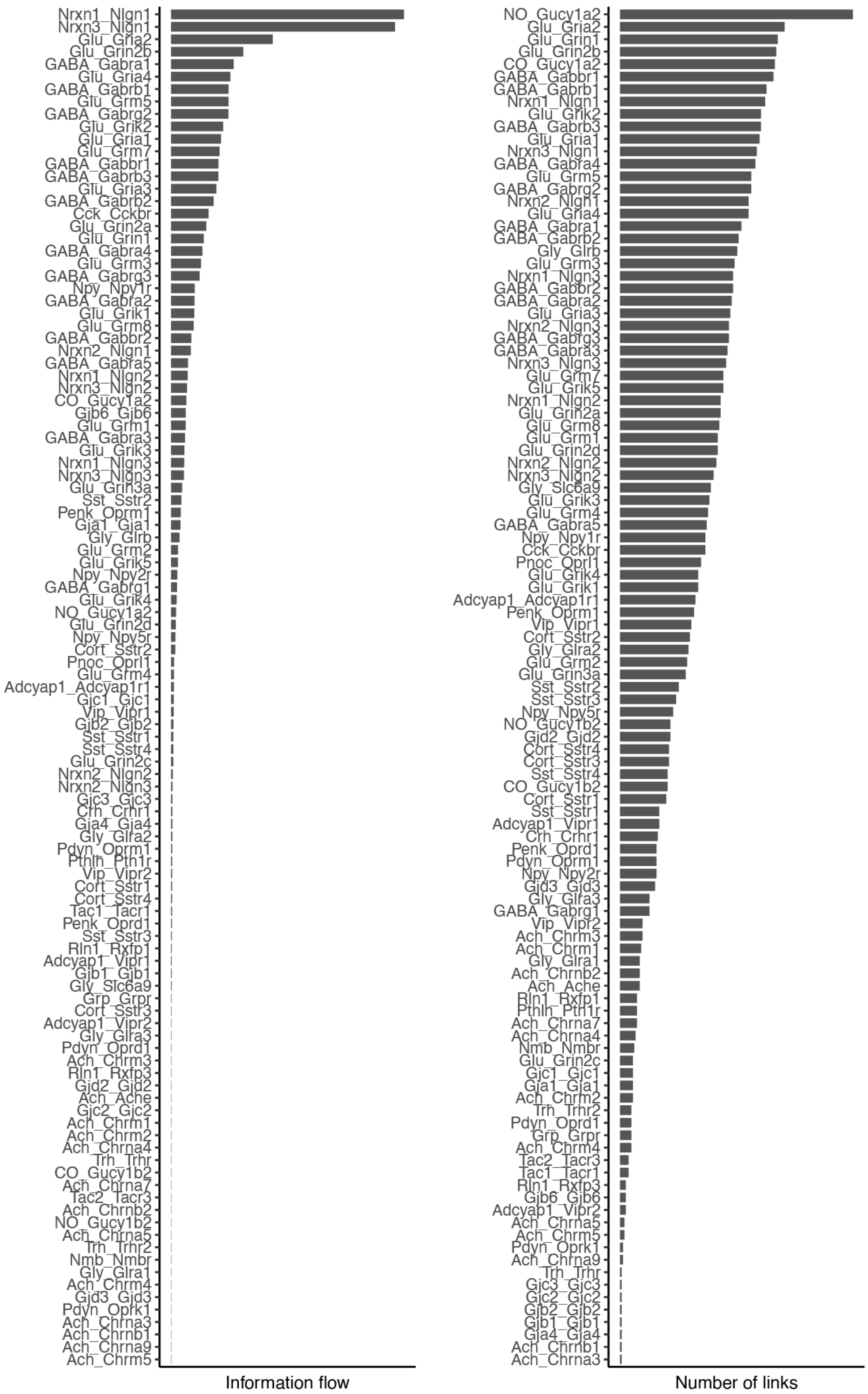


**Supplementary Figure 10. Bar charts illustrating the information flow (left) and number of links (right) for detected interaction pairs for intercellular communication networks of mouse primary visual cortex.**

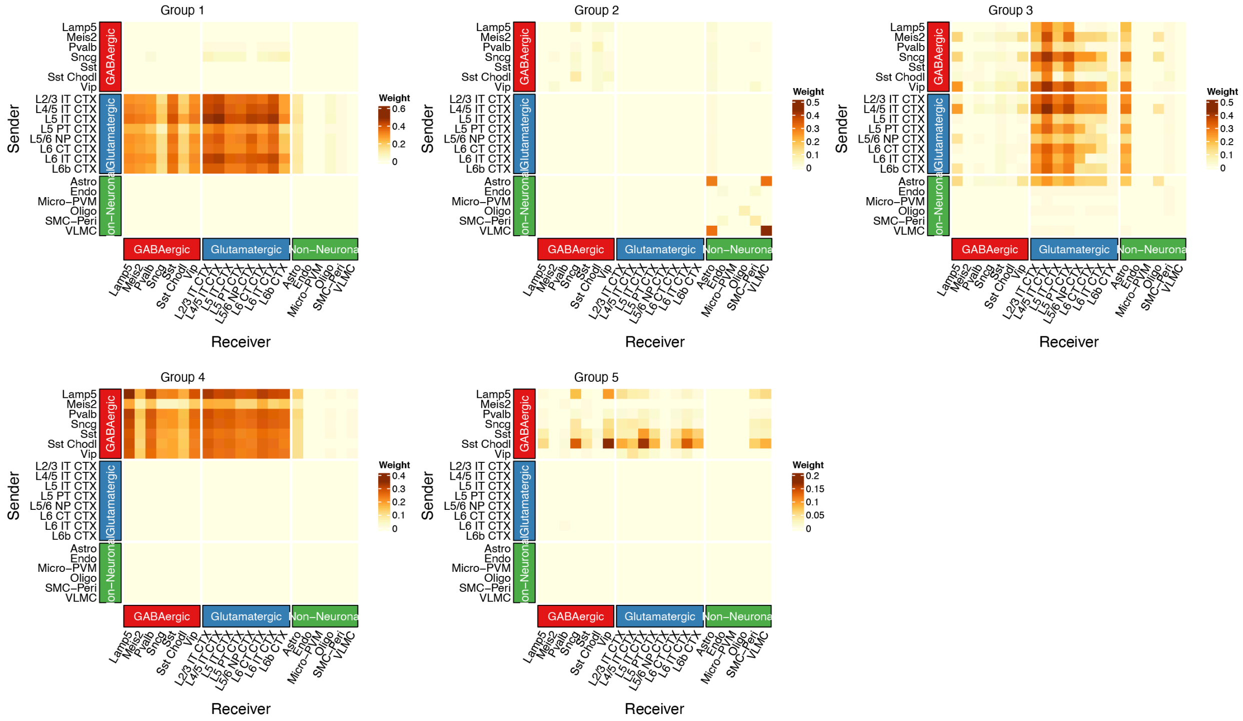


**Supplementary Figure 11. Heatmaps of the aggregated communication networks for the five interaction groups in Figure 4c.**

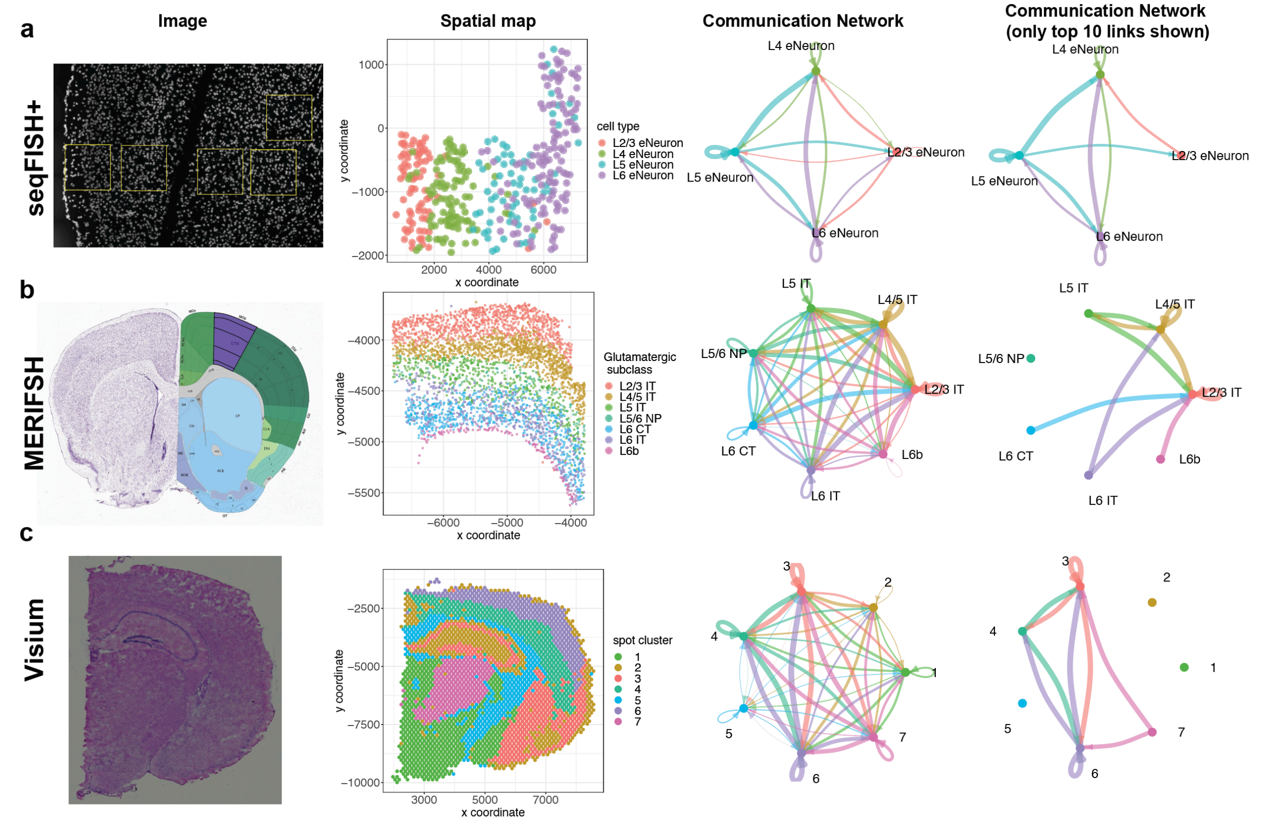


**Supplementary Figure 12. Inference of communication network for three spatial transcriptomics datasets.**

(a-c) The raw tissue slice image/brain region annotation diagram (first panel), spatial map (second panel), full aggregated communication network (third panel), and aggregated communication network with top 10 links shown (fourth panel), for three spatial transcriptomics datasets generated by different techniques including seqFISH+ (a), MERFISH (b), and Visium (c). In spatial maps, a dot represents the centroid of a cell (for a-b) or a spot (for c). The communication network summarizes the communication strength over interaction pairs (i.e., the "weight" aggregation method). For seqFISH+ and Visium datasets, the communication networks are directly calculated from the spatial transcriptomics; for the MERFISH dataset, the communication network is calculated using single-cell RNA-seq data (4,461 cells of seven glutamatergic subclasses) for the same brain region. For (b), the brain region annotation image (Image credit: Allen Institute for Brain Science. [http://atlas.brain-map.org/atlas?atlas=1#atlas=1&plate=100960348]) highlights the mouse primary motor cortex region (in purple); the spatial map only shows the glutamatergic cells in one representative coronal slice (slice id: mouse1_slice212).

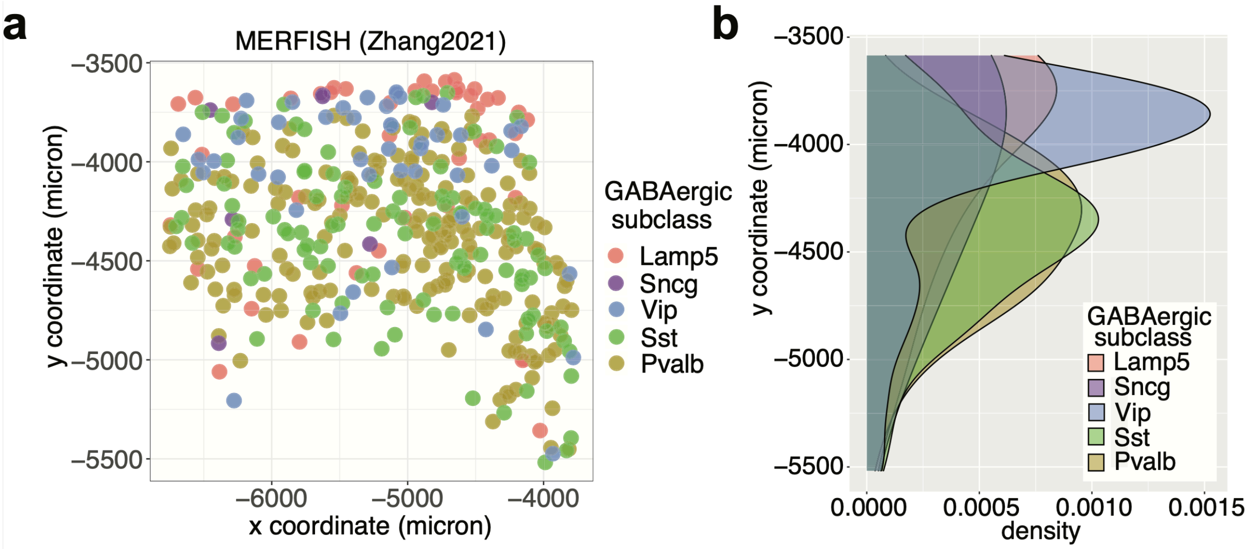


**Supplementary Figure 13. The spatial map of GABAergic neurons for the MERFISH dataset.**

(a) Spatial map of five subclasses of GABAergic neurons in a coronal slice (slice id: mouse1_slice212). A dot represents the centroid of a cell.

(b) Distribution of the y-axis coordinate for the five GABAergic subclasses shown in (a). The y-axis coordinate roughly represents the range from cortical layer L1 (top) to L6b (bottom).
